# A large sensory and multi-omics evaluation unraveled chemical and genetic basis of orange flavor

**DOI:** 10.1101/2023.07.10.548426

**Authors:** Zhen Fan, Kristen A. Jeffries, Xiuxiu Sun, Gabriela Olmedo, Wei Zhao, Matthew R. Mattia, Ed Stover, John A. Manthey, Elizabeth A. Baldwin, Seonghee Lee, Frederick G. Gmitter, Anne Plotto, Jinhe Bai

**Affiliations:** Horticultural Sciences Department, IFAS Gulf Coast Research and Education Center, University of Florida, Wimauma, FL 33598; Horticultural Research Laboratory, USDA-ARS, Fort Pierce, FL, 34945; Daniel K. Inouye U.S. Pacific Basin Agricultural Research Center, USDA-ARS, Hilo, Hawaii 96720; Horticultural Sciences Department, IFAS Citrus Research and Education Center, University of Florida, Lake Alfred, FL 33850

## Abstract

Sweet orange (*Citrus sinensis*) exhibits limited genetic diversity and high susceptibility to Huanglongbing (HLB). New HLB-tolerant orange-like hybrids are promising alternatives. However, the genetic control of key flavor compounds in oranges remains unknown. Evaluating 179 juice samples, including oranges, mandarins, *Poncirus trifoliata* and hybrids, distinct volatile compositions were found. A random forest model predicted untrained samples with 78% accuracy and identified 26 compounds crucial for orange flavor. Notably, seven esters—methyl hexanoate, ethyl hexanoate, ethyl 3-hydroxyhexanoate, ethyl octanoate, methyl butanoate, ethyl butanoate, and ethyl 2-methylbutanoate—differentiated orange from mandarin. Cluster analysis showed six esters with shared genetic control. Differential gene expression analysis identified *CsAAT1*, an *alcohol acyltransferase* responsible for ester production in orange. Its activity was validated through overexpression assays. A SNP-based DNA marker in the CDS region accurately predicted phenotypes. This study enhances our understanding of orange flavor compounds, their biosynthetic pathways, and expands breeding options for orange-like cultivars.

## Introduction

Citrus production in the United States has been devastated by Huanglongbing (HLB) or citrus greening disease, especially in Florida where historically citrus production was 90% sweet orange (*Citrus sinensis* (L.) Osbeck), a citrus type highly susceptible to *Candidatus* Liberibacter asiaticus, the bacterium considered responsible for the disease. In the 2022-23 season, Florida citrus production had the lowest output in nine decades, taking acreage down to ∼50% and production to ∼10% of levels prior to the arrival of the disease ^1^. Thus far, there is no effective way to control HLB. The production costs have increased dramatically due to additional disease management practices and more frequent insecticide applications, which only slow down the disease progression but does not cure infected trees ^2^. By 2017, the cumulative economic loss due to HLB was estimated to be $6 billion ^3^. The most sustainable solution to maintain orange production is to plant cultivars that are resistant or tolerant to HLB.

Currently, only a narrow range of citrus cultivars are classified as sweet orange, *C. sinensis*, which are all derived from cumulative somatic mutations derived from a presumed single complex interspecific introgression hybrid of mandarin (*Citrus reticulata* Blanco.) and pummelo (*Citrus maxima* (Burm.) Merr.) ^4,5^. Therefore, the limited genetic diversity of sweet orange has rendered it difficult to find a resistant source within the species. Breeding new cultivars via outcrossing with HLB tolerant materials provides the best breeding solution. *Poncirus trifoliata* (L.), conferring tolerance to HLB, has been used for citrus breeding. It is challenging to improve the fruit quality of hybrids harboring *Poncirus* introgressions, because fruits of the parent *P. trifoliata* have unacceptable flavor, and the hybrids with *Citrus* present various levels of off-flavor even after a few generations of backcrossing to elite citrus cultivars ^6,7^. However, after generations of backcrosses to mandarin, new generations of orange-like hybrids are available, such as ‘US SunDragon’ included in this study, which has *P. trifoliata* in its pedigree and low *P. trifoliata* off-taste ^3^. ‘US SunDragon’ exhibits tolerance to HLB and a high resemblance to orange flavor, and thus is a promising new variety to sustain the orange juice industry in Florida.

One key to accelerating the development of orange-like hybrids and enhancing the acceptance of using non-*C. sinensis* in orange juice is to define fruit components important for the distinctive flavor and aroma characteristics in orange. Studies before 2005 focused on the identification of abundant and odor-active compounds in orange fruit or juice samples alone. The fruity aroma esters ethyl 2-methylpropanoate, ethyl butanoate and ethyl 2-methylbutanoate, 3a,4,5,7a-tetrahydro-3,6-dimethyl-2(3H)-benzofuranone (“wine lactone”) and the grassy aroma (*Z*)-hex-3-enal were found to be the most potent volatiles in Valencia and Navel oranges, based on aroma extract dilution analysis (AEDA) and odor activity values (OAVs) ^8,9^. Among those, ethyl butanoate decreased dramatically during juice pasteurization ^10^. Recently, efforts shifted to comparing aroma profiles between orange and other citrus species to find alternatives for HLB-sensitive *C. sinensis*. Using the same techniques, but comparing ‘Valencia’ orange to a mandarin cultivar, a recent study identified ethyl butanoate, ethyl 2-methylbutanoate, octanal, decanal, and acetaldehyde as the most odor-active volatiles in orange ^11^. Nevertheless, all above-mentioned studies are of small scale, consisting of three cultivars at most. A new chemical and sensory evaluation, testing a diverse collection of cultivars within and across citrus species was desired to validate and expand on the previous results.

Woody perennial plants like citrus have a long juvenile phase and require a long and expensive evaluation process before releasing new hybrid cultivars. To date, only one genetic map of fruit aroma was created in citrus, using a biparental population between two mandarin cultivars ^12^. A total of 206 quantitative trait loci (QTLs) were identified for 94 volatiles, demonstrating the abundance of natural variation in the mandarin volatilome. However, no aroma genes were directly discovered and validated via traditional forward-genetic approaches. On the other hand, using reverse-genetic approaches, various terpene synthases (TPSs) in citrus were identified and functionally validated ^13,14^. TPSs with different enzymatic activities catalyze production of monoterpenes and sesquiterpenes found in citrus fruit. *CsTPS1*, discovered in *C. sinensis*, is responsible for valencene production in orange ^13^. A natural InDel in the promoter region of *CsTPS1* was associated with deficiency of valencene production in some mandarin cultivars ^15^. Two transcription factors, CitERF71 and CitAP2.10, regulate *E*-geraniol and valencene production by interaction with *CitTPS16* and *CsTPS1*, respectively ^16,17^. Nevertheless, genetic studies on citrus fruit flavor are limited to the terpene biosynthesis pathway. No information about other aroma biosynthetic genes or molecular markers in citrus is available.

Here, we report a comprehensive chemical and flavor evaluation over 179 harvest/accession combinations, including sweet orange (*C. sinensis*), mandarin (*C. reticulata*), poncirus (*P. trifoliata*) and citrus hybrids with poncirus introgressions (hereafter poncirus hybrids) with the objective to predict orange flavor using chemical data and to deduce the most important and key chemical compounds contributing to orange flavor. Integration of chemical clustering and transcriptome analyses facilitated the identification of a novel *alcohol acyltransferase* (*CsAAT1*), catalyzing the biosynthesis of both straight- and branched-chain esters contributing to orange flavor. Its biological function was validated in citrus fruits using overexpression transient assays. Last, a new DNA marker based on an exonic SNP was developed and tested.

## Results

### Volatilome and flavor quality of mandarin (*C. reticulata*), sweet orange (*C. sinensis*), *Poncirus trifoliata* and poncirus introgressed hybrids

A total of 60 volatiles, consisting of five alcohols, eight aldehydes, twelve esters, two ketones, fourteen monoterpenes, one terpene ester, one terpene ketone, four terpene alcohols, and thirteen sesquiterpenes (Table S4) were identified and quantified for 198 harvest/accession combinations (Data S1). Based on PCA of the volatilome, clear separation was observed among poncirus (*P. trifoliata*), sweet orange (*C. sinensis*) and mandarin (*C. reticulata*) (Fig. 1A). However, poncirus hybrids exhibited a wider distribution, overlapping with all three species on the first two principal components (PCs) (Fig. 1B), reflecting their admixture genetics. Some poncirus hybrids showed similar volatile profiles to those of orange, revealed by their proximity to orange on the first two PCs (Fig. 1B) and clustering close to four orange cultivars in the hierarchical clustering analysis (Fig. 1C). In general, compared to orange, two poncirus cultivars were higher in ketones, terpene ester, sesquiterpenes and terpene alcohols, while lower in alcohols, aldehydes, esters and terpene ketone (Fig. 2A). After generations of modified backcrossing, poncirus hybrids recovered levels similar to orange, for a majority of volatiles, except for esters (Fig. 2A). Specifically, significantly higher concentrations of methyl butanoate (*p* = 1.6E-13, Bonferroni corrected), ethyl butanoate (*p* = 2.5E-10), ethyl hexanoate (*p* = 1.3E-5), ethyl 3-hydroxyhexanoate (*p* = 3.9E-18) and ethyl octanoate (*p* = 4.1E-10) were detected in orange than in poncirus hybrids (Table S5). In addition to chemical measurements, juice samples were also evaluated for orange and mandarin flavor attributes by a trained panel. Similar to volatile data, sensory scores of orange flavor for poncirus hybrids exhibited large variation, ranging from 1.1 to 8.1 (Fig. 2B, Data S2.xlsx). One hybrid, ‘US SunDragon’ (SUN), scored consistently high for orange flavor (orange score > average orange score for orange cultivars, 6.4) across 11 out of 14 harvests (Fig. 2B), while its volatile profile was not the closest to orange (Fig. 1C). These results indicate volatiles found in orange, did not equally contribute to the orange flavor, with some having more importance than others.

**Figure 1.**
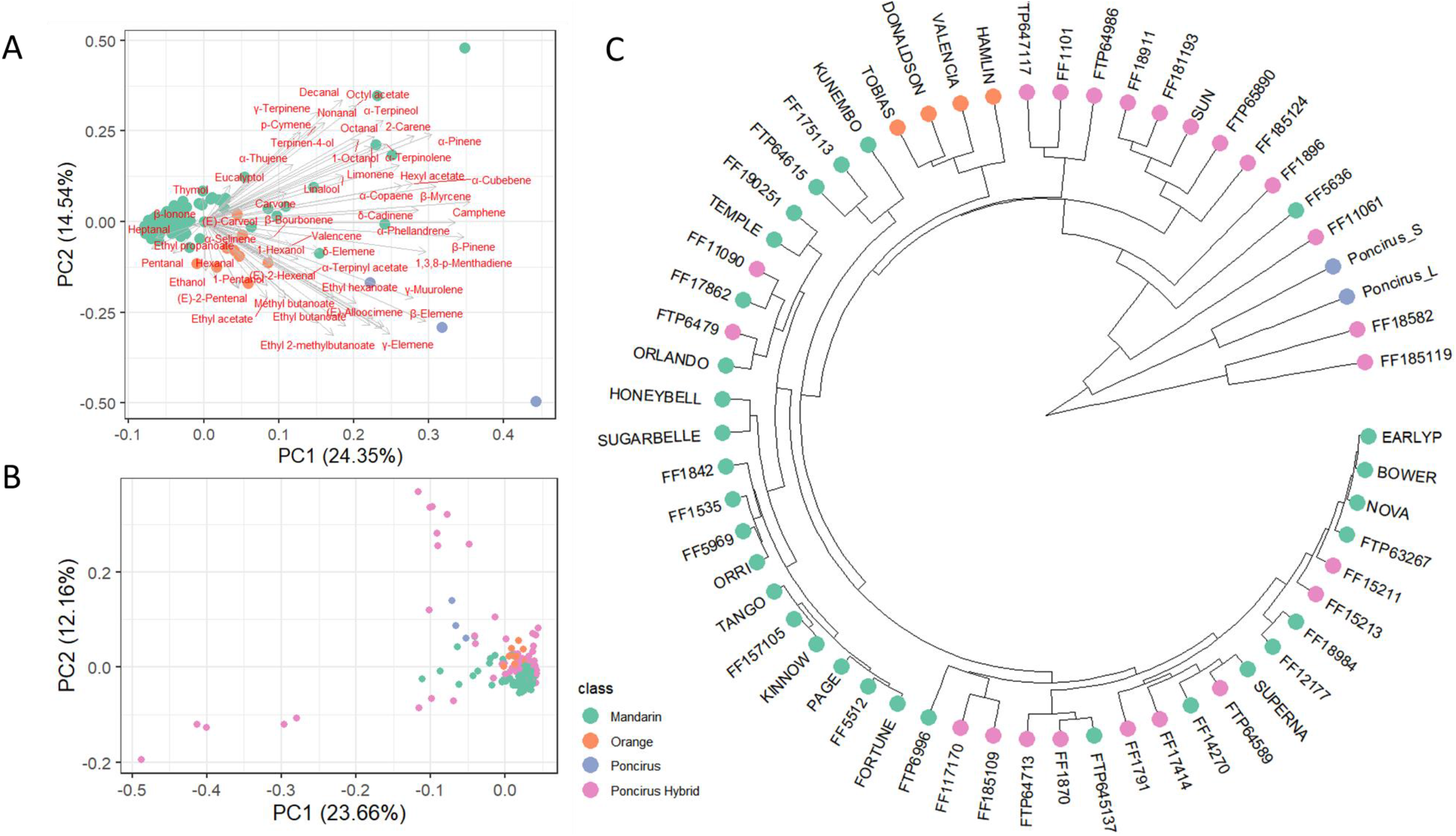
(A) Biplot of first two principal components based on volatile data. Individual hybrids are colored according to their breeding class. Samples of hybrids, predominately mandarin (*Citrus reticulata*), sweet orange (*Citrus sinensis*), and *Poncirus trifoliata* are included in this principal component analysis (PCA). (B) *Poncirus* introgressed hybrids were added to PCA. (C) Hierarchical clustering of all cultivars, constructed using absolute distance matrix of volatile data.

**Figure 2.**
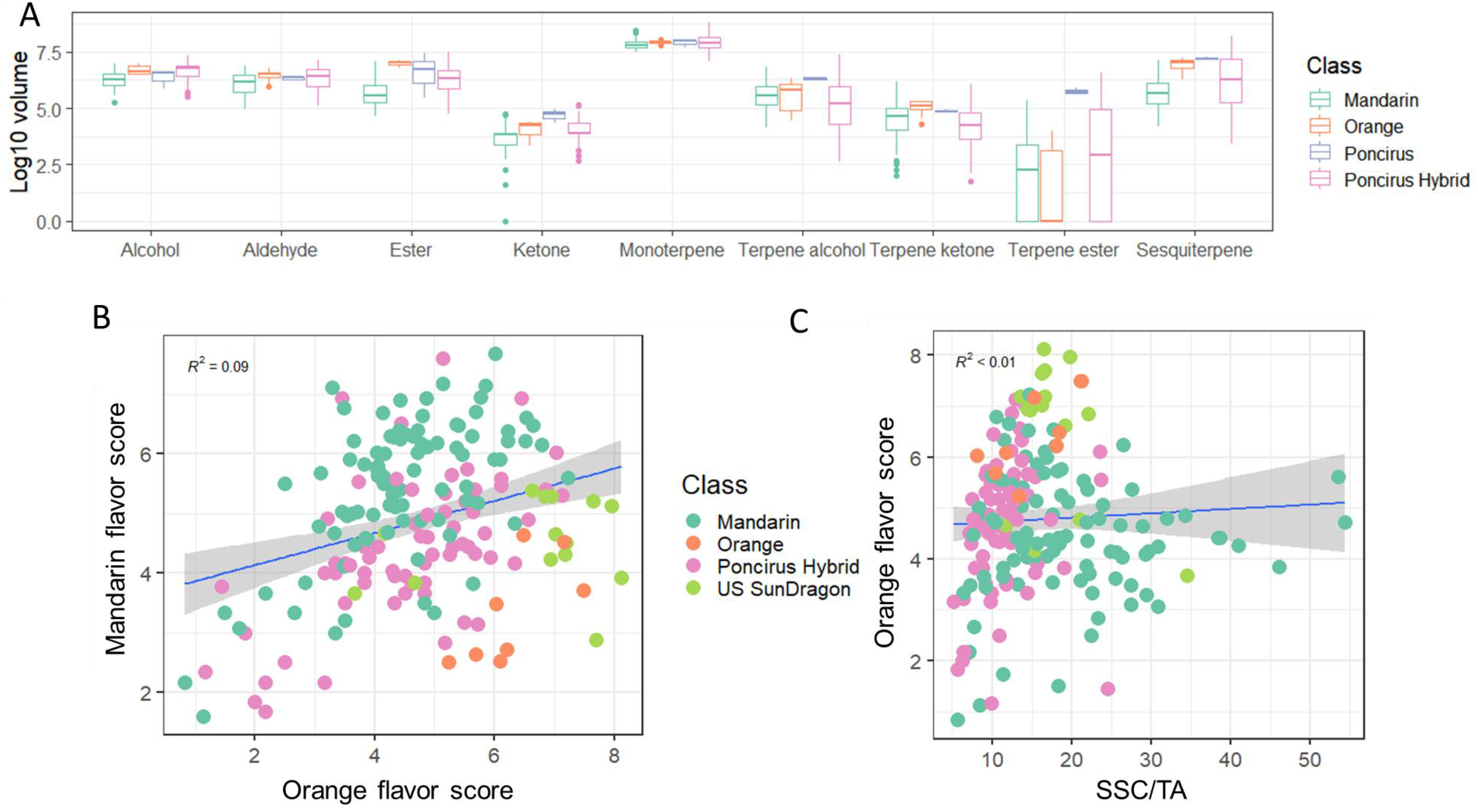
(A) Comparisons of total volume of volatile classes among different species. (B) Mandarin flavor scores regressed against orange flavor scores. Samples of ‘US SunDragon’ are colored in light green. (C) Orange flavor scores regressed against SSC/TA ratio.

### Prediction of orange flavor using machine learning (ML) algorithms

Unlike mandarin flavor (R^2^ = 0.21, Fig. S1), orange flavor (R^2^ < 0.01, Fig. 2C) was poorly predicted by SSC/TA. Therefore, to predict orange flavor entails an integration of sugars, acids, bitter compounds, and volatiles. Here, we tested six popular statistical models ^18,19^ for predictions of orange, mandarin, and relative orange (orange score – mandarin score) flavors, based on measurements of SSC, TA, pH, limonin, nomilin and 60 volatiles. To test the prediction accuracy of our models in real-world scenarios, all samples (n = 19) collected in the 2021-2022 season were not included in the training, but only used for testing. The random forest (RF) model consistently performed the best across three flavor attributes and two testing schemes (Fig. 3), with R^2^ of 0.67, 0.74 and 0.78 for respective orange, mandarin, and relative orange flavors for the 2021-22 samples (Fig. 3B). The high predictive ability of the RF model gave us high confidence that RF model could be used to identify important chemical compounds and predict orange flavor in future seasons.

**Figure 3.**
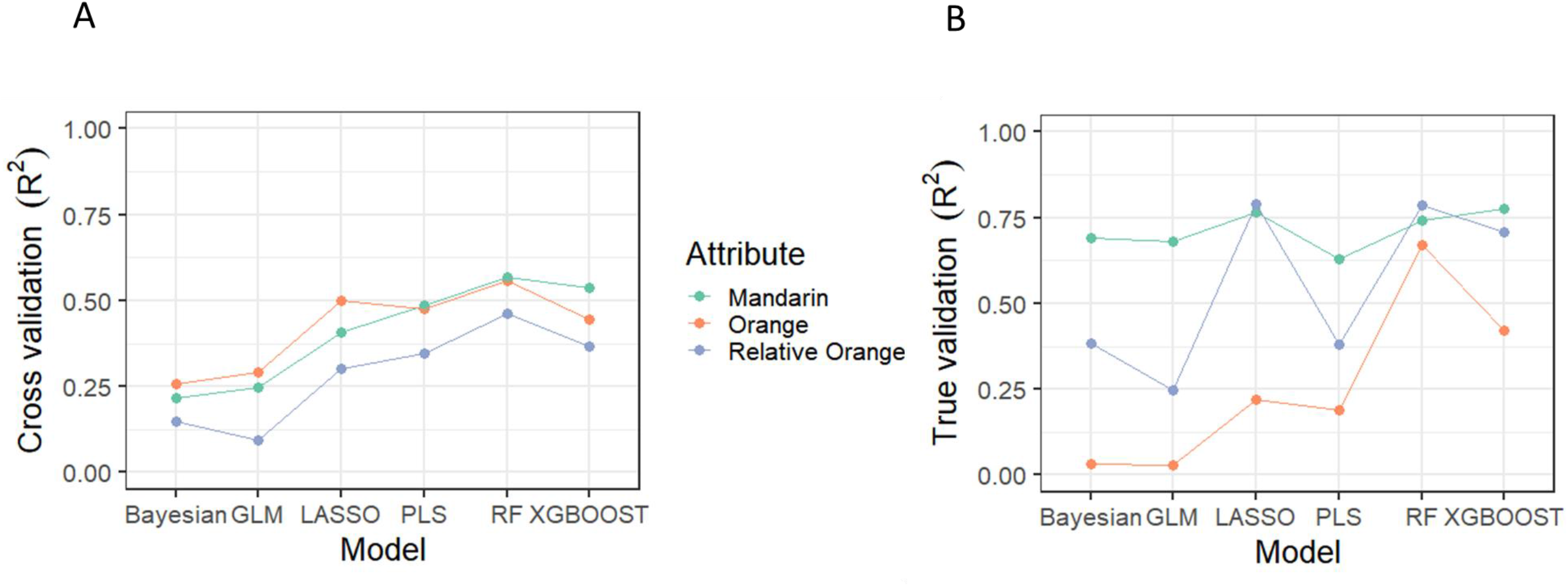
Six prediction models (Bayesian linear model (Bayesian), General Linear Mixed model (GLM), LASSO regression, Partial Least Squares (PLS), Random Forest (RF), Extreme Gradient Boosting decision tree (XGBoost)) used to predict orange, mandarin and relative orange flavors, and tested with the 10-fold cross validation (A); true validation dataset which includes 19 samples collected from season 2021-2022 (B). Y axis is the R^2^ value.

### Important compounds for predicting orange flavor

Among SSC, TA, pH, limonin, nomilin and 60 volatiles, RF models identified 26 chemical compounds, important in prediction of relative orange and orange flavors. The important compounds included eight esters, two aliphatic alcohols, two aliphatic aldehydes, one aliphatic ketone, seven terpenes or terpene derivatives, both limonoids, SSC, TA, pH and SSC/TA (Table 1). Both bitter limonoids, limonin and nomilin, exhibit negative effects on relative orange flavor, while both positive and negative effects were observed for terpenes and terpene derivatives. Two alcohols (1-pentanol and 1-hexanol) and two aldehydes ((*E*)-2-pentenal and (*E*)-2-hexenal) showed positive relationships with relative orange flavor. 1-Octen-3-one, which was reported to be the most odor-active aliphatic ketone in orange juice ^20,21^, showed high positive effect on relative orange flavor. A negative effect of SSC/TA was observed for relative orange flavor, consistent with the observation of lower SSC/TA in orange compared to mandarin in previous studies ^22^. Seven esters (methyl hexanoate, ethyl hexanoate, ethyl 3-hydroxyhexanoate, ethyl octanoate, methyl butanoate, ethyl butanoate, and ethyl 2-methylbutanoate), all with positive effect on orange flavor, were the only compounds shared between the prediction model for orange flavor scores and the classification model for orange (Table S6). The total level of seven esters explained 21% percent of the total variation in relative orange flavor (Fig. S2). A specific hybrid with *P. trifoliata* in its background, ‘US SunDragon’, had high orange flavor as well as high total esters (Fig. S3). On the other hand, esters were negative predictors for mandarin flavor, yet only three contributors, SSC/TA, (*E*)-2-pentenal and SSC showed positive effects on mandarin flavor. In summary, orange flavor requires high amounts of esters, which are the key and essential compounds, as well as several alcohols and aldehydes, but low limonoids, lower SSC/TA than mandarin and a balanced combination of terpenes.

**Table 1.** Important compounds (Variable Importance in Projection (VIP) > 6) for prediction of orange and mandarin flavor in citrus fruit, and their relative intensity (relative orange = orange-mandarin)

| Compound | Class | Relative orange |  |  | Orange |  |  | Mandarin |  |  |
| --- | --- | --- | --- | --- | --- | --- | --- | --- | --- | --- |
|  |  | VIP | Effect | Rank | VIP | Effect | Rank | VIP | Effect | Rank |
| Methyl hexanoate* | Ester | 63.87 | P | 1 |  |  |  |  |  |  |
| Ethyl hexanoate* | Ester | 34.01 | P | 2 |  |  |  | 10.33 | N | 5 |
| pH |  | 27.83 | N | 3 | 7.64 | N | 13 |  |  |  |
| 1-Pentanol | Alcohol | 24.54 | P | 4 |  |  |  |  |  |  |
| Limonin | Limonoid | 24.24 | N | 5 | 8.89 | N | 9 |  |  |  |
| SSC/TA |  | 21.82 | N | 6 | 10.50 | P | 8 | 31.32 | P | 2 |
| 1-Octen-3-one | Ketone | 17.88 | P | 7 |  |  |  |  |  |  |
| α-Cubebene | Sesquiterpene | 16.86 | P | 8 | 35.85 | P | 2 | | | |
| (E)-2-Pentenal | Aldehyde | 13.18 | P | 9 |  |  |  | 8.07 | P | 6 |
| (E)-2-Hexenal | Aldehyde | 12.96 | P | 10 | 8.65 | P | 10 |  |  |  |
| Ethyl 3-hydroxyhexanoate* | Ester | 11.92 | P | 11 |  |  |  | 55.38 | N | 1 |
| Ethyl octanoate* | Ester | 10.57 | P | 12 |  |  |  |  |  |  |
| TA |  | 10.23 | P | 13 |  |  |  | 7.80 | N | 7 |
| Nomilin | Limonoid | 8.19 | N | 14 | 61.73 | N | 1 |  |  |  |
| α-Terpinyl acetate | Terpene ester | 7.62 | P | 15 | | | | | | |
| Methyl butanoate* | Ester | 7.30 | P | 16 |  |  |  |  |  |  |
| Ethyl butanoate* | Ester | 7.08 | P | 17 |  |  |  |  |  |  |
| Ethyl 2-methylbutanoate* | Ester | 6.79 | P | 18 |  |  |  |  |  |  |
| SSC |  |  |  |  | 10.99 | P | 5 | 14.86 | P | 3 |
| β-Bourbonene | Sesquiterpene | | | | | | | 13.90 | N | 4 |
| Carvone | Terpene ketone |  |  |  | 18.41 | P | 3 | 7.64 | N | 8 |
| Caryophyllene | Sesquiterpene |  |  |  |  |  |  | 7.36 | N | 9 |
| β-Elemene | Sesquiterpene | | | | | | | 6.11 | N | 10 |
| Ethyl propanoate | Ester |  |  |  | 11.75 | P | 4 |  |  |  |
| Humulene | Sesquiterpene |  |  |  | 10.84 | N | 6 |  |  |  |
| Terpinen-4-ol | Terpene alcohol |  |  |  | 10.78 | N | 7 |  |  |  |
| 1-Hexanol | Alcohol |  |  |  | 8.47 | P | 11 |  |  |  |
| (E)-Alloocimene | Monoterpene |  |  |  | 7.67 | N | 12 |  |  |  |
| δ-Cadinene | Sesquiterpene | | | | 6.67 | P | 14 | | | |

\*Overlap with important predictors for classification as orange based on pedigree data

### Flavor compounds and their related transcriptome networks in *Citrus*

Chemical clustering showed 50 out of 66 analyzed compounds were clustered into nine groups (Fig. 4), consistent with their chemical classes. There were two groups for sesquiterpenes (ST1, ST2), two for monoterpenes (MT1, MT2), one for alcohol (ALC1), two for aldehydes (ALD1, ALD2) and two for esters (E1, E2). Compounds within a cluster might share the same genetic control and indicate existing natural genetic variation across different cultivars. To identify gene co-expression modules that jointly altered with chemical groups, the weighted correlation network analysis (WGCNA, Fig. S4) was built using RNAseq data from juice samples of eleven accessions with different volatile profiles. Six gene co-expression modules (Fig. S5A, Data S3) were correlated to chemical clusters, including E2 to MEivory (r = 0.77), E1 to MEdarkolivegreen (r = 0.82), TA to MEindianred4 (r = 0.67), SSC to MEbrown4 (r = 0.80), ALD1 to MEplum2 (r = 0.87), and ST2 to MEfirebrick4 (r = 0.82, only observed in dendrogram with individual compounds, Fig. S5B). *CsTPS1* (Cs4g_pb015710) was within the co-expression module MEfirebrick4, found positively correlated with concentrations of five sesquiterpenes (valencene, γ-muurolene, α-selinene, β-elemene and humulene), consistent with the previous functional characterization ^13^. The identification of *CsTPS1* validated the effectiveness of WGCNA to discover metabolite biosynthetic genes. Otherwise, the SSC-related module MEbrown4 contained 26 genes, enriched for galactose metabolic process (GO:0006012). The ALD1-related module MEplum2 contained a D-isomer specific 2-hydroxyacid dehydrogenase (Cs1g_pb014450), a fatty acid desaturase (Cs6g_pb001810), and an alpha/beta hydrolase (Cs9g_pb018430), all potentially involved in fatty acid metabolic process. Lastly, a GDSL lipase/esterase like gene (Cs3g_pb021240) in the module MEivory was related with E2 concentrations.

**Figure 4.**
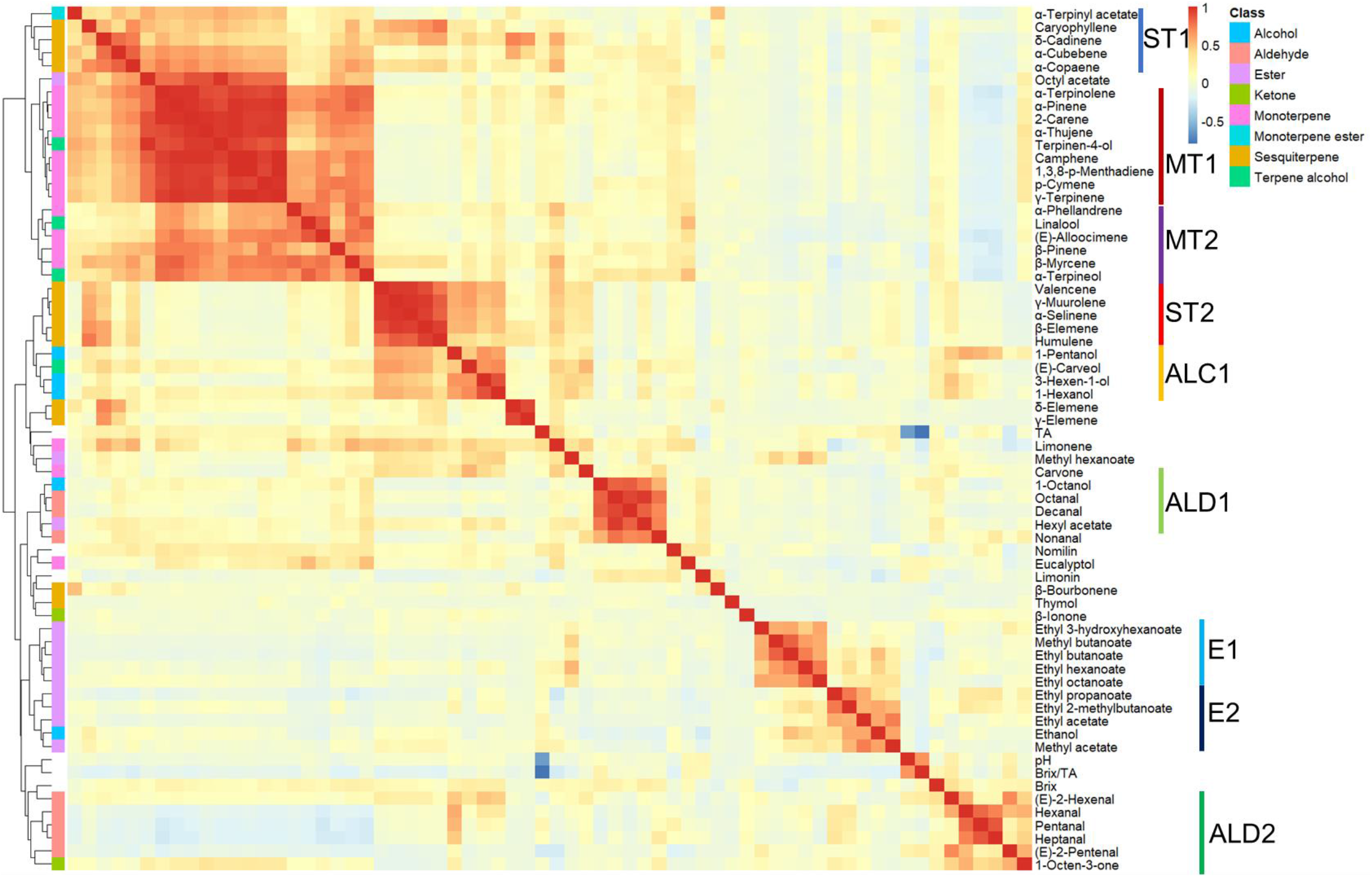
Heatmap of correlations among chemical abundances. Each cell is colored according to Pearson correlation coefficient (−1 to 1). Hierarchical clustering (HCA) is shown on the left side of the plot. Nine groups (right side of the plot) are assigned based on chemical class and HCA. The letters in group names represent chemical classes: sesquiterpenes (ST), monoterpenes (MT), alcohols (ALC), aldehydes (ALD), esters (E).

### Discovery of *CsAAT1* responsible for ester biosynthesis in orange

Using chemical clustering, five of seven key esters for orange flavor were clustered into E1 group and methyl hexanoate showed high correlation with E1 group despite not being clustered in E1 (Fig. 4). To identify candidate genes responsible for ester production in orange, we compared gene expression between five ester producers (all esters in E1 group) and six non-producers (Table S1). After filtering with FDR adjusted *p*-value < 0.05 and fold change > 2, only 93 upregulated genes (0.2% of total genes) were identified. Among them, a novel *alcohol acyltransferase* Cs6g_pb013840.1 (*CsAAT1*) showed four-fold increase of expression in ester producers than in non-producers (Fig. 5A), with a correlation coefficient of 0.74 with the first eigenvector of E1 cluster (Fig. 5B). The expression of *CsAAT1* was ripening-induced, as an increase of expression was observed along the ripening process while no expression was observed in roots, leaves, or fruits 45 days after flowering (Fig. S6) ^23^. A close examination of synteny regions from multiple genome assemblies revealed that the *CsAAT1* and its flanking regions were incomplete in the *C. sinensis* genome V2 (Cs6g_pb013840) ^24^, and complete but not annotated in the genome V3 ^4^. After reannotation with RNAseq data and homolog search, a tandem array of two *AATs* was identified, with an interspace of 8593 bp (Fig. 5D). However, only *CsAAT1* was expressed across samples, while *CsAAT1t* (the tandem homolog of *CsAAT1*) had close to zero expression (Fig. 5D). Sanger sequencing of cDNA confirmed that *CsAAT1* was 1347 bp long, with only one exon, encoding a polypeptide of 449 amino acid residues and a predicted molecular mass of 50.56 kDa. It contained characteristic HXXXDG (residues 155–160), and DFGWG (residues 378–382) motifs of the BAHD protein family (Fig. S7) ^25^. In a maximum likelihood tree with 23 diverse acetyltransferases, *CsAAT1* was clustered within the clade including three strawberry alcohol acetyltransferases (*FvAAT, FcAAT1* and *FvAAT*) and *RhAAT* identified in *Rosa hybrida* (Fig. 5C), with sequence similarity between 39.3% and 40.9% to four AATs. The sequence similarity to strawberry AATs explained the resemblance of ester profile to cultivated strawberry (*Fragaria ×ananassa*) as all esters in orange except for ethyl 3-hydroxyhexanoate could be detected in strawberry ^18,26,27^. Alignment among *CsAAT1* homologs in *C. sinensis* (orange), *C. maxima* (pummelo), *C. reticulata* (mandarin) and *P. trifoliata* revealed four non-synonymous polymorphisms unique in orange and an insertion of Tyrosine at residue 205, only found in *C. maxima* and *C. sinensis* (Fig. S7), indicating the functional allele of *CsAAT1* was likely inherited from pummelo, consistent with the previous admixture results ^5^.

**Figure 5.**
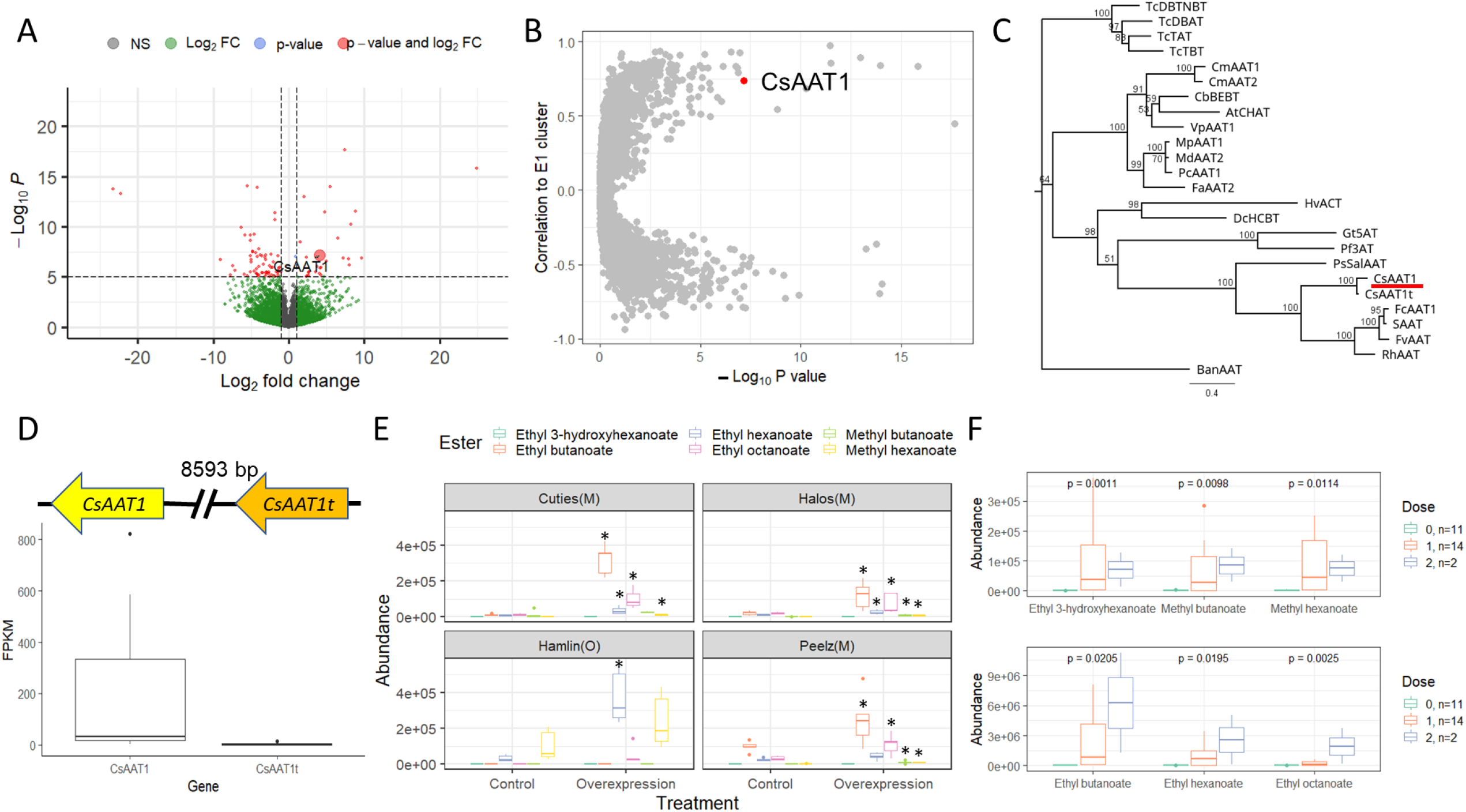
(A) Volcano plot of log_2_ fold change against -log_10_ P-value comparing ester producers to non-producers. *CsAAT1* is highlighted with a large dot. (B) Correlation of expression of *CsAAT1* (red dot) to the first eigenvector of E1 cluster. X axis is -log10 P-value. Y axis is correlation to first eigenvector of E1 cluster. Individual dots represent genes. (C) Maximum likelihood tree of 24 acyltransferases with 100 fast bootstrapping. *CsAAT1* is underlined. Branches are annotated with bootstrapping values. (D) Comparison of expression of *CsAAT1* and its tandem replicate *CsAAT1t*. (E) Comparisons of production for six esters between control fruits and fruits agroinfiltrated with *CsAAT1* overexpression construct across four cultivars. M indicates mandarin cultivar. O indicates orange cultivar. Asterisks indicate P-values < 0.05 based on Student t tests. (F) Concentrations of six esters with 0,1,2 dosages of the orange allele of *CsAAT1* across 27 cultivars. The number of samples in each category is provided. P-values from Kruskal-Wallis tests are provided.

### Functional validation and DNA marker for *CsAAT1*

The function of *CsAAT1* was validated via transient overexpression assays in three mandarin cultivars (ester non-or low-producer) at the fully ripe stage, and one orange cultivar at the small green stage (1 to 2cm in diameter). Across mandarin cultivars, fruits agroinfiltrated with the overexpression construct of *CsAAT1* had significantly increased production of methyl butanoate, methyl hexanoate, ethyl butanoate, ethyl hexanoate, and ethyl octanoate (Fig. 5E, Table S7). The highest concentrations were found in three ethyl esters, likely facilitated by a high volume of free substrate, ethanol, in the ripe citrus fruits (Data S1). Although there was no increase of ethyl 3-hydroxyhexanoate in the overexpressed fruits, significant increases of the other four branched-chain esters were observed, especially ethyl trans-2-butenoate (Fig. S8), suggesting *CsAAT1* was also capable of synthesizing branched-chain esters. On the other hand, transient overexpression of *CsAAT1* in young green fruits of orange resulted in higher production of ethyl hexanoate, methyl hexanoate and ethyl octanoate (Fig. 5E). The difference in ester profile comparing to overexpressed mandarin may be related to changing substrate availability during fruit development. A high-resolution melting (HRM) marker was developed to target 1122A->C. It showed 100% sensitivity and 85.1% accuracy to predict the production of six esters (Fig. 5F, Fig. S9, Table S3). All accessions with zero dose of the *CsAAT1* orange allele (n=11) produced no esters in fruit. Four cultivars with heterozygous calls produced low levels of esters (Table S3), indicating potential additional genetic controls for ester production.

## Discussion

Unique odor of different citrus types is the result of a distinct composition of volatiles. Despite characteristic compounds found in some citrus such as grapefruit ^28^, no consensus of key compounds for orange has been reached. It was generally accepted that a combination of twenty to thirty volatiles is needed to imitate orange flavor ^20^. However, this hypothesis was challenged by other studies. Using aroma reconstitution experiments, a mix of fourteen aroma active compounds in their actual concentrations highly resembled orange juice reconstituted from concentrate ^21^. Feng et al. cut the list down to five key compounds which were ethyl butanoate, ethyl 2-methylbutanoate, octanal, decanal, and acetaldehyde ^11^. Reactive aldehydes such as acetaldehyde, (*Z*)-hex-3-enal, neral, and geranial, imparting a pleasant green and citrus note to fresh squeezed orange juices, are present at supra threshold levels in fresh squeezed oranges, yet diminish in processed orange juice ^20^. The two aldehydes, (*E*)-2-pentenal and (*E*)-2-hexenal, found to contribute to orange flavor in our work, fit into this category. However, concentrations of straight-chain aldehydes such as octanal, nonanal, and decanal were not found important for orange flavor in our study because both decanal and octanal are two major aldehydes in other citrus species such as mandarins ^29–31^. Our prediction models, incorporating the largest sensory and chemical dataset of citrus cultivars to date, identified seven esters, including the most odor active compounds in orange juice, ethyl 2-methylbutanoate and ethyl butanoate ^9,11^, as key compounds for orange flavor. Consistent with these results, orange juice made from HLB affected orange fruits, with a lower concentration of esters, was often described as lacking orange flavor ^32^.

Since citrus types produce a large diversity and quantity of terpenes, genetic research has focused on the discovery of genes for terpene biosynthesis up to now. Some characterized terpene biosynthetic genes include limonene synthase ^33^, linalool synthase ^34^, γ-terpinene synthase ^35^, sabinene synthase ^36^ and valencene synthase ^13^. Less is known about genetic controls for other aroma active compounds in citrus such as esters. *Alcohol acyltransferase (AAT)* is a member of the BAHD family and is able to accept a range of alcohol and acyl-CoA substrates to form esters ^25^. *AATs* have been functionally characterized in a variety of fruit crops, such as strawberry ^37^, melon ^38^ and banana ^39^, among others. Our phylogeny revealed that the novel *CsAAT1* belonged to Clade III according to previous classification ^25^, together with other *AATs*. More specifically, *CsAAT1* is most phylogenetically related to three AATs in strawberry. Strawberry AATs can utilize short- and medium-chain, branched, and aromatic acyl-CoA and alcohol molecules as substrates ^27,37^ to produce a variety of straight- and branched-chain esters including six key esters found in orange. The function prediction of *CsAAT1* via phylogenetic analysis is congruent with results from fruit overexpression assays and marker tests. Ester biosynthesis is also regulated by the availability of substrates, which may depend on the activity of various catabolic pathways (e.g., lipid breakdown) ^37^, which could explain our observation of an increased amount of hexanoic acid esters, but not butanoic acid esters in small green overexpressed oranges. A previous study has shown orange was distinguished from other citrus cultivars by the lowest levels of oleic and palmitoleic, and the highest levels of linolenic and arachidic acids ^40^. Future study is needed to explore genetic variation in the early steps of ester biosynthesis among citrus germplasm, as well as fatty acid availability during fruit development.

Current juice regulations limit adding fruits from mandarin hybrids not classified as *Citrus sinensis* to 10% to maintain the “orange juice” label, leaving the citrus juice industry vulnerable to disease epidemics such as HLB when growing oranges in a monoculture ^3^. Our findings show that some poncirus-introgressed hybrids such as ‘US Sundragon’ and FF11061 are organoleptically similar to orange, more HLB tolerant than orange, and can potentially be used in orange juice, widening selections that can substitute for traditional orange cultivars. Using a large collection of accessions from multiple citrus types, we determined esters were the key compounds for perceiving orange flavor and differentiating orange from other citrus types. We discovered a novel alcohol acyltransferase, *CsAAT1*, which catalyzes the production of both straight- and branched chain esters in orange fruit. Our work will greatly accelerate the breeding of orange-like hybrids. With the aid of the new DNA marker for orange flavor, seedlings can be screened at an early stage, long before they produce fruits, and orange flavor can be more rapidly recovered with fewer generations of backcrossing.

## Materials and methods

### Fruit sampling

Fruits were sampled from mature trees grown at the A.H. Whitmore Citrus Research Foundation Farm, Groveland, FL (28.687502, -81.886090) or at the USDA/ARS Research Farm, Fort Pierce, FL (27.433215, - 80.427057). Trees were self-rooted for hybrids or grafted on rootstocks for named cultivars. Hybrids were selected based on tree health and general quality, spanning a large range in shapes, colors, seed numbers, and flavors, especially during the first season (2016-2017) ^41^. Hybrids that produced fruit that were too sour or bitter in the first two years were eliminated from further observations. Hybrids of interest (healthy trees and acceptable flavor) were harvested multiple times at 2-to-4-week intervals to determine optimum maturity. Fruits were then taken to the USDA/ARS laboratory in Fort Pierce, FL, washed, sanitized and manually juiced using a reamer-type juicer (Oster Model 3183, Household Appliance Sales and Service, Niles, IL, USA, or Vinci™ Hand Free Juicer, Vinci® Housewares, La Mirada, CA, USA). Fruits were split into four batches of equal number of pieces to account for four replications. Aliquots were taken for measurements of soluble solids concentration (SSC), titratable acidity (TA), limonin, nomilin, and volatiles. A minimum of one liter was used for sensory evaluation. All juice samples were stored at -20 °C until analysis. A total of 198 harvest/accession combinations were analyzed for volatile production, and/or evaluated for orange and mandarin flavors, with 178 samples having both evaluations. Information about sample ID, harvest time, harvest location and pedigree for individual samples is in Data S2.

### Flavor evaluation

Most panelists had been trained and practiced descriptive sensory evaluation of citrus juices for over 10 years. In this study, ten to twelve of these panelists rated citrus hybrids for orange and mandarin flavors using a linear scale with numeral anchors from 0 to 15, and word anchors at 1 = “low”, 7.5 = “medium”, and 15 = “high”. A reference standard was provided for each of these descriptors at each session.

Unpasteurized orange juice locally produced (Al’s Family Farm®, Fort Pierce, FL) and tangerine juice (“gourmet pasteurized” from Natalie’s Orchid Island Juice Company (Fort Pierce, FL)) were the references for orange and mandarin flavor, respectively. Orange and mandarin juice standards were given an intensity value of 12 on the 0-15 scale, as they were considered the representative flavor for those descriptors. Panelists met each year for five or six sessions to practice and refresh their memories about descriptors. They tasted samples that were evaluated from the previous year and that represented typical orange and mandarin flavors, then discussed their evaluations. The panel leader gathered and averaged their ratings from the discussion and entered the values using the Feedback Calibration Method (FCM®) feature in Compusense Cloud (Compusense^®^, Guelph, ON, Canada). Panelists then returned to the tasting booths and evaluated the same samples presented in coded cups, using the FCM® method. Samples were served as 45 mL in 118 mL (4-oz) plastic soufflé cups (Solo^®^ Cups Co., Urbana, IL) at 14 °C, in a randomized order across panelists. Panelists tasted up to 40 samples per season, with no more than 4 samples per day, repeated in two daily sessions.

### Volatile identification and quantification

Volatiles were analyzed using a headspace-solid phase microextraction (SPME) - gas chromatography-mass spectrometry system as previously reported ^42^. Briefly, 6 mL of juice were sealed in 20-mL vials and stored at -20 °C until analysis. The juice sample was incubated at 40 °C for 30 min and then a 2 cm tri-phase SPME fiber (50/30 μm DVB/Carboxen/ PDMS, Supelco, Bellefonte, PA, USA) was inserted to the headspace to collect and concentrate volatiles for 30 min. The SPME fiber was then inserted into the injector of an Agilent 7890 GC coupled with a 5975 MS detector (Agilent Technologies, Palo Alto, CA, USA) for 15 min at 250 °C. The column was a DB-5 (60 m x 0.25 mm i.d., 1.00 μm film thickness, J&W Scientific, Folsom, CA, USA). Mass units were monitored from 30 to 250 m/z and ionized at 70 eV. Volatile identification and quantification of peak areas were conducted with MassHunter Workstation software (Version 10.0; Agilent Technologies). Initial identification was done by mass spectra searches with the NIST library (Version 14, match score > 0.9). The identification was then confirmed by comparing the retention indices generated by running standard C6-C17 alkane mixture under the same conditions as the samples with online resources (NIST Chemistry WebBook and Flavornet.org). Each sample had four replicates.

### Soluble solids concentration, titratable acidity and limonoids quantification

Juice was centrifuged and soluble solids concentration (SSC), titratable acidity (TA) and limonoids were measured using the supernatant. SSC was determined by a digital refractometer (Atago RX-5000cx, Tokyo, Japan). TA was measured by titration of 10 mL of supernatant with 0.1 mol L^−1^ sodium hydroxide (NaOH) to the final pH of 8.1 using a titrator (Dosino model 800, Metrohm, Herisau, Switzerland). Quantification of limonin and nomilin was performed via Liquid chromatography-tandem mass spectrometry (LC-MS/MS) with a 1290 Infinity II UPLC coupled with a 6470 triple quadrupole MS (Agilent, Santa Clara, CA, USA). The MS was operated in MRM mode and nomilin was detected with a *m/z* 515.3 precursor ion and a *m/z* 411.2 product ion with fragmentor and collision energy voltages set to 135V and 14V respectively. Limonin was detected with a *m/z* 471.2 precursor ion and a *m/z* 425.2 product ion with fragmentor and collision energy voltages set to 135V and 19V respectively. Quantification of limonoids was performed by integrating the area under the chromatographic peak and calculating the amount of each compound based on standard curves (R^2^ ≥ 0.99 with range 0.006 - 25 mg L^-1^). Each sample had four replicates that were averaged.

### Prediction models and statistical analyses

Model training and testing were conducted using tidymodels package in R ^43^. A total of 160 samples evaluated from 2016 to 2021 was used to train models. Seven prediction models including Generalized Linear Model (GLM), Least Absolute Shrinkage and Selection Operator Model (LASSO), Bayesian Model, Partial Least Square Model (PLS), XGBoost Model and Random Forest Model (RF) were used to predict scores for orange, relative orange (orange score – mandarin score) and mandarin flavors. Two hyperparameters were tuned for the PLS model including the maximum proportion of original predictors and the number of PLS components to retain. Three hyperparameters were tuned for XGBoost and RF including the number of predictors that randomly sampled at each split, the number of trees contained in the ensemble, and the minimum number of data points in a node required for the node to be split further. Tuning for hyperparameters was conducted using a grid searching method and the best values were determined using 10-fold cross validation. To evaluate the performance of the final models, we used two testing datasets, a 10-fold cross validation set, and a true validation set including 19 samples collected in the 2021-22 season. Since RF performed the best across both validation sets, it was used to infer important compounds for prediction of flavor attributes. Variable Importance in Projection (VIP) scores were extracted from RF models using VIP package in R. Compounds with VIP > 6 were identified as important compounds. Principal component analysis (PCA) and hierarchical clustering of chemical compounds were constructed with absolute distance matrix using prcomp and dist functions in R.

### RNA extraction, sequencing, and alignment

RNA was extracted from fruit juice samples of 11 accessions (Table S1) having distinct volatile profiles. RNeasy Plant Mini kits (QIAGEN Co., Hilden, Germany) were used for extraction. Illumina (San Diego, CA, USA) 150-bp pair-end sequencing was performed on the Illumina NovoSeq platform by Novogene Co. (Sacramento, CA, USA). On average, 7.1 Gb data was obtained for each sample. Reads were aligned to sweet orange genome V2 ^24^ using STAR aligner with the default parameters ^44^. Reads for each transcript were counted using htseq-count ^45^ in Union mode excluding non-unique reads.

### Weighted correlation network analysis (WGCNA) and Differential gene expression (DGE) analysis

Weighted correlation network for gene expression was conducted using WGCNA package in R ^46^. Transcript counts for 11 accessions were converted into Fragments Per Kilobase of transcript per Million mapped reads (FPKM). The input transcripts were filtered with total FPKM across 11 samples > 1, and coefficient of variation > 0.58, a total of 4092 transcripts remained for the analysis. The value of softPower was set to 8, chosen by the “pickSoftThreshold” function. The minimum module size was set to 10 and deepsplit parameter was set to 2. The rest of the parameters in WGCNA were set to default. To find relationships between co-expression modules and chemical compounds, two correlation matrices were built. The first one included the first eigenvectors of all co-expression modules and abundances of all chemical compounds; the second one consisted of the first eigenvectors of all co-expression modules and chemical clusters. The distance matrix was computed as 1 – correlation matrix. Hierarchical clustering analysis was performed to determine relationships between co-expression modules and chemical compounds. Differentially expressed genes (DEGs) comparing between ester producers (n = 5) and non-producers (n = 6) were identified using DESeq2 package in R ^47^. The input list of transcripts was pre-filtered using only those with total counts > 10. The final list of DEGs was determined with FDR adjusted p-value < 0.05 and fold change > 2.

### Sanger sequencing of *CsAAT*1 cDNA, gene tree and high-melting resolution marker

Total RNA of ‘Valencia’ orange juice was used to synthesize cDNA. To amplify the full length of *CsAAT1* cDNA, two primers (CsAAT1_1F: ATGGAAATTGGCATTGTCTCAAGAG, CsAAT1_1,340R: TCAGAATCGACAAACGTCAAACAATA) were designed according to the curated coding region of *CsAAT1* based on evidence of genome sequences of SWOv2 and SWOv3 ^4^ and RNAseq alignment. cDNA of *CsAAT1* was amplified with touchdown PCR and a gel confirmed the size of amplicon. Sanger sequencing for unpurified PCR product was conducted at Azenta Inc. using two primers (CsAAT1_1,340R, CsAAT1_658R: TGTACTTGGCTTCTCTGAACCA). The sequencing result confirmed the CDS sequence of *CsAAT1* (NCBI accession: OR003937). To build a gene tree of acyltransferases including *CsAAT1*, protein sequences of 23 acyltransferases including alcohol acyltransferases involved in the biosynthesis of esters from strawberry ^37^, melon ^38^, banana ^39^ and apple ^48^ were downloaded from NCBI (Table S2). Protein sequences were aligned using Clustal Omega V1.2.2. A maximum likelihood tree with 100 fast bootstrapping was built with GAMMA GTR model using RAxML. A high-resolution melting marker (HRM, CsAAT1_1,087F: GCCTTGAAATTTTCCAGTTGGG, CsAAT1_1,175R: TCTCCAAAAATGCCGCTCCA) was designed to target the SNP 1122A->C. PCR and melting curve genotyping was conducted according to previous studies ^49^. The HRM marker was tested for 27 cultivars which had volatile data from multiple harvests (Table S3). Sensitivity was calculated as 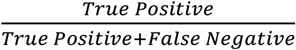 and accuracy was measured as 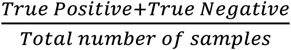.

### Transient fruit overexpression assay

The full-length cDNA of *CsAAT1* flanked by attB sequences was synthesized by Twist Bioscience Inc. (San Francisco, CA, USA). The target sequence was first introduced into pDONR/Zeo vector (ThermoFisher, Inc. Waltham, MA, USA) via Gateway cloning, and then inserted into overexpression vector pMDC32 ^50^. The empty vector pMDC32 and overexpression vector pMDC32::*CsAAT1* were transformed into *Agrobacterium tumefaciens* strain EHA105. Transformation to citrus fruits was performed according to Shen et al. ^17^. Fruits from three commercial mandarin brands (‘Peelz™’, ‘Cuties®’, ‘Halos®’) were purchased at a local grocery store and small green fruits of ‘Hamlin’ orange (1 to 2 cm in diameter), in which esters should not be produced based on tissue-specific expression results (Fig. S6), were harvested from the research farm. Two treatments were performed on separate fruits. Five biological replicates with one fruit per rep were performed for each treatment. Four injections with 3ml Agrobacterium suspension each were made for mandarin fruits. For orange fruits, the injection stopped when the whole fruit was wet. Pulp tissue (whole fruit for orange) was collected after 7 days of incubation in dark at room temperature. All samples were analyzed with the GC-MS with the aforementioned conditions.

## Supplementary figure legends

Figure S1. Mandarin flavor scores regressed against SSC/TA. Individual dots are colored according to their breeding class. The best fitted linear model is plotted with 0.95 confidence intervals. R squared value is annotated on the top left.

Figure S2. Relationship between relative orange flavor scores (orange flavor – mandarin flavor) and total volume of seven esters. Individual dots are colored according to their breeding class. The best fitted linear model is plotted with 0.95 confidence intervals. R squared value is annotated on the top left.

Figure S3. The total abundance of seven esters (top) and relative orange score (bottom) are plotted for each accession. Bars and dots are colored according to their breeding class. Error bars are calculated based on multiple harvests.

Figure S4. Co-expression modules assigned using weighted correlation network analysis (WGCNA). The top plot shows hierarchical clustering results. The bottom track shows the assigned module based on the dynamic tree cutting algorithm.

Figure S5. Dendrogram of correlations among eigengenes (the first eigenvector) of co-expression modules and the first eigenvectors of chemical clusters (A). Dendrogram of correlations among eigengenes of co-expression modules and individual compounds (B).

Figure S6. Expression of *CsAAT1* across five different tissues. 45DAF and 142DAF : fruit harvested 45 and 142 days after flowering, respectively.. A190DAF: fruits harvested after 190 days after flowering. Raw data were downloaded from public RNAseq results deposited at CPBD (http://citrus.hzau.edu.cn/index.php). Four replicates were used for each tissue.

Figure S7. Protein alignment among homologs of *CsAAT1* in *Poncirus trifoliata* (Pt6g009140.1 & Pt6g009150.1), *Citrus reticulata* (Cre6g_014390.1 & Cre6g_014400.1) and *Citrus maxima* (Cg_ZPY6g_009600.1 & Cg_ZPY6g_009610.1) with *CsAAT1* and *CsAAT1t*.

Figure S8. Comparisons of production for four branched-chain esters between control citrus fruits and fruits agroinfiltrated with *CsAAT1* overexpression construct. M indicates mandarin cultivar. O indicates orange cultivar. Asterisks indicate P-values < 0.05 based on Student t tests.

Figure S9. Melting curve patterns of samples with different dosages of the orange allele of *CsAAT1*.

## Supplementary table legends

Table S1. RNAseq sample information

Table S2. Information of acyltransferases used in the gene tree.

Table S3. HRM marker genotype and average ester abundance across multiple harvests for 28 cultivars.

Table S4. Volatile compounds identified and quantified for citrus samples.

Table S5. Contrasting individual volatile between groups.

Table S6. Variable Importance in Projection (VIP) scores in classification model for sweet orange.

Table S7. Comparison of ester abundances between overexpressed and control samples.

## Supplementary data legends

Data S1. Raw data of volatile abundances.

Data S2. Median sensory and chemical measurements across replicates of each harvest/accession combination.

Data S3. Genes contained in each cluster of the weighted correlation network.

## Data availability

The full-length cDNA of *CsAAT1* is available at NCBI, with accession number OR003937. Raw RNAseq data are available on NCBI under BioProject accession no. PRJNA977630.

## Contributions

Z.F., E.B., E.S., A.P. and J.B. conceptualized the project. Z.F., K.J. G.O., A.P. and J.B. developed the methodology. Z.F., K.J., X.S., G.O., W.Z., M.M., J.M., A.P. and J.B. curated the data, conducted the formal analysis and investigation, generated the data visualizations and wrote the original draft of the manuscript. E.S., E.B., S.L., F.G., A.P. and J.B. reviewed and edited the manuscript.

## Citations

1. USDA - National Agricultural Statistics Service - Florida - Citrus Statistics. https://www.nass.usda.gov/Statistics_by_State/Florida/Publications/Citrus/Citrus_Statistics/.

2. Ariel Singerman. FE1006/FE1006: Cost of Production for Processed Oranges Grown in Central Florida (Ridge), 2015/16. https://edis.ifas.ufl.edu/publication/FE1006 (2020).

3. Stover, E. et al. Rationale for reconsidering current regulations restricting use of hybrids in orange juice. Horticulture Research 2020 7:1 7, 1–7 (2020).

4. Wang, L. et al. Somatic variations led to the selection of acidic and acidless orange cultivars. Nature Plants 2021 7:7 7, 954–965 (2021).

5. Wu, G. A. et al. Genomics of the origin and evolution of Citrus. Nature 2018 554:7692 554, 311–316 (2018).

6. Deterre, S. et al. Secondary metabolite composition in citrus × Poncirus trifoliata hybrids. Proceedings of the Florida State Horticultural Society 126, 206–215 (2013).

7. Deterre, S. C. et al. Effect of Poncirus trifoliata on the chemical composition of fruits in pedigrees of Citrus scion hybrids. Sci Hortic 277, 109816 (2021).

8. Hinterholzer, A. & Schieberle, P. Identification of the Most Odour-active Volatiles in Fresh, Hand-extracted Juice of Valencia Late Oranges by Odour Dilution Techniques. Ltd. Flavour Fragr. J 13, 49–55 (1998).

9. Buettner, A. & Schieberle, P. Evaluation of aroma differences between hand-squeezed juices from Valencia late and Navel oranges by quantitation of key odorants and flavor reconstitution experiments. J Agric Food Chem 49, 2387–2394 (2001).

10. Nisperos-Carríedo, M. O. & Shaw, P. E. Comparison of Volatile Flavor Components in Fresh and Processed Orange Juices. J Agric Food Chem 38, 1048–1052 (1990).

11. Feng, S., Suh, J. H., Gmitter, F. G. & Wang, Y. Differentiation between Flavors of Sweet Orange (Citrus sinensis) and Mandarin (Citrus reticulata). J Agric Food Chem 66, 203–211 (2018).

12. Yu, Y. et al. Identification of QTLs controlling aroma volatiles using a ‘Fortune’ x ‘Murcott’ (Citrus reticulata) population. BMC Genomics 18, 1–16 (2017).

13. Sharon-Asa, L. et al. Citrus fruit flavor and aroma biosynthesis: isolation, functional characterization, and developmental regulation of Cstps1, a key gene in the production of the sesquiterpene aroma compound valencene. The Plant Journal 36, 664–674 (2003).

14. Alquézar, B., Rodríguez, A., de la Peña, M. & Peña, L. Genomic analysis of terpene synthase family and functional characterization of seven sesquiterpene synthases from citrus sinensis. Front Plant Sci 8, 1481 (2017).

15. Yu, Q. et al. Deficiency of valencene in mandarin hybrids is associated with a deletion in the promoter region of the valencene synthase gene. BMC Plant Biol 19, 1–11 (2019).

16. Li, X. et al. Transcription factor CitERF71 activates the terpene synthase gene CitTPS16 involved in the synthesis of E-geraniol in sweet orange fruit. J Exp Bot 68, 4929–4938 (2017).

17. Shen, S. L. et al. CitAP2.10 activation of the terpene synthase CsTPS1 is associated with the synthesis of (+)-valencene in ‘Newhall’ orange. J Exp Bot 67, 4105–4115 (2016).

18. Fan, Z. et al. Strawberry sweetness and consumer preference are enhanced by specific volatile compounds. Hortic Res 8, 1–15 (2021).

19. Colantonio, V. et al. Metabolomic selection for enhanced fruit flavor. Proc Natl Acad Sci U S A 119, e2115865119 (2022).

20. Ruiz Perez-Cacho, P. & Rouseff, R. Processing and Storage Effects on Orange Juice Aroma: A Review. J Agric Food Chem 56, 9785–9796 (2008).

21. Averbeck, M. & Schieberle, P. H. Characterisation of the key aroma compounds in a freshly reconstituted orange juice from concentrate. European Food Research and Technology 229, 611–622 (2009).

22. Porat, R., Deterre, S., Giampaoli, P. & Plotto, A. The flavor of citrus fruit. Biotechnology in Flavor Production 1–31 (2016) doi:10.1002/9781118354056.CH1.

23. Liu, H. et al. Citrus Pan-Genome to Breeding Database (CPBD): A comprehensive genome database for citrus breeding. Mol Plant 15, 1503–1505 (2022).

24. Xu, Q. et al. The draft genome of sweet orange (Citrus sinensis). Nature Genetics 2012 45:1 45, 59–66 (2012).

25. D’Auria, J. C. Acyltransferases in plants: a good time to be BAHD. Curr Opin Plant Biol 9, 331–340 (2006).

26. Ulrich, D., Kecke, S. & Olbricht, K. What do we know about the chemistry of strawberry aroma? J Agric Food Chem 66, 3291–3301 (2018).

27. González, M. et al. Aroma development during ripening of Fragaria chiloensis fruit and participation of an alcohol acyltransferase (FcAAT1) gene. J Agric Food Chem 57, 9123–9132 (2009).

28. Buettner, A. & Schieberle, P. Evaluation of key aroma compounds in hand-squeezed grapefruit juice (Citrus paradisi Macfayden) by quantitation and flavor reconstitution experiments. J Agric Food Chem 49, 1358–1363 (2001).

29. Huang, M. et al. Characterization of the Major Aroma-Active Compounds in Peel Oil of an HLB-Tolerant Mandarin Hybrid Using Aroma Extraction Dilution Analysis and Gas Chromatography-Mass Spectrometry/Olfactometry. Chemosens Percept 10, 161–169 (2017).

30. Obenland, D., Collin, S., Sievert, J. & Arpaia, M. L. Mandarin flavor and aroma volatile composition are strongly influenced by holding temperature. Postharvest Biol Technol 82, 6–14 (2013).

31. Moshonas, M. G. & Shaw, P. E. Quantitation of Volatile Constituents in Mandarin Juices and Its Use for Comparison with Orange Juices by Multivariate Analysis. J Agric Food Chem 45, 3968–3972 (1997).

32. Dala-Paula, B. M. et al. Effect of huanglongbing or greening disease on orange juice quality, a review. Front Plant Sci 9, 1976 (2019).

33. Morehouse, B. R. et al. Functional and Structural Characterization of a (+)-Limonene Synthase from Citrus sinensis. Biochemistry 56, 1706–1715 (2017).

34. Shimada, T. et al. Characterization of three linalool synthase genes from Citrus unshiu Marc. and analysis of linalool-mediated resistance against Xanthomonas citri subsp. citri and Penicilium italicum in citrus leaves and fruits. Plant Science 229, 154–166 (2014).

35. Suzuki, Y. et al. Characterization of γ-terpinene synthase from Citrus unshiu (Satsuma mandarin). BioFactors 21, 79–82 (2004).

36. Kohzaki, K. et al. Characterization of a sabinene synthase gene from rough lemon (Citrus jambhiri). J Plant Physiol 166, 1700–1704 (2009).

37. Aharoni, A. et al. Identification of the SAAT gene involved in strawberry flavor biogenesis by use of DNA microarrays. Plant Cell 12, 647–662 (2000).

38. El-Sharkawy, I. et al. Functional characterization of a melon alcohol acyl-transferase gene family involved in the biosynthesis of ester volatiles. Identification of the crucial role of a threonine residue for enzyme activity. Plant Mol Biol 59, 345–362 (2005).

39. Beekwilder, J. et al. Functional Characterization of Enzymes Forming Volatile Esters from Strawberry and Banana. Plant Physiol 135, 1865–1878 (2004).

40. Moufida, S. & Marzouk, B. Biochemical characterization of blood orange, sweet orange, lemon, bergamot and bitter orange. Phytochemistry 62, 1283–1289 (2003).

41. Baldwin, E. et al. Handling & Processing Section Accelerating Implementation of Huanglongbing-tolerant Hybrids As New Commercial Cultivars for Fresh and Processed Citrus. Proc. Fla. State Hort. Soc 132, 177–181 (2019).

42. Bai, J., Baldwin, E., Hearn, J., Driggers, R. & Stover, E. Volatile profile comparison of USDA sweet orange-like hybrids versus ‘Hamlin’and ‘Ambersweet’. HortScience 49, 1262–1267 (2014).

43. Max, K. H. W. Tidymodels: a collection of packages for modeling and machine learning using tidyverse principles. https://www.tidymodels.org.

44. Dobin, A. et al. STAR: ultrafast universal RNA-seq aligner. Bioinformatics 29, 15–21 (2013).

45. Anders, S., Pyl, P. T. & Huber, W. HTSeq—a Python framework to work with high-throughput sequencing data. Bioinformatics 31, 166–169 (2015).

46. Langfelder, P. & Horvath, S. WGCNA: An R package for weighted correlation network analysis. BMC Bioinformatics 9, 1–13 (2008).

47. Love, M. I., Huber, W. & Anders, S. Moderated estimation of fold change and dispersion for RNA-seq data with DESeq2. Genome Biol 15, 1–21 (2014).

48. Souleyre, E. J. F., Greenwood, D. R., Friel, E. N., Karunairetnam, S. & Newcomb, R. D. An alcohol acyl transferase from apple (cv. Royal Gala), MpAAT1, produces esters involved in apple fruit flavor. FEBS J 272, 3132–3144 (2005).

49. Oh, Y. et al. Genomic Characterization of the Fruity Aroma Gene, FaFAD1, Reveals a Gene Dosage Effect on γ-Decalactone Production in Strawberry (Fragaria × ananassa). Front Plant Sci 12, 639345 (2021).

50. Curtis, M. D. & Grossniklaus, U. A Gateway Cloning Vector Set for High-Throughput Functional Analysis of Genes in Planta. Plant Physiol 133, 462 (2003).

