## Supplementary Figures for "A large sensory and multi-omics evaluation unraveled chemical and genetic basis of orange flavor"

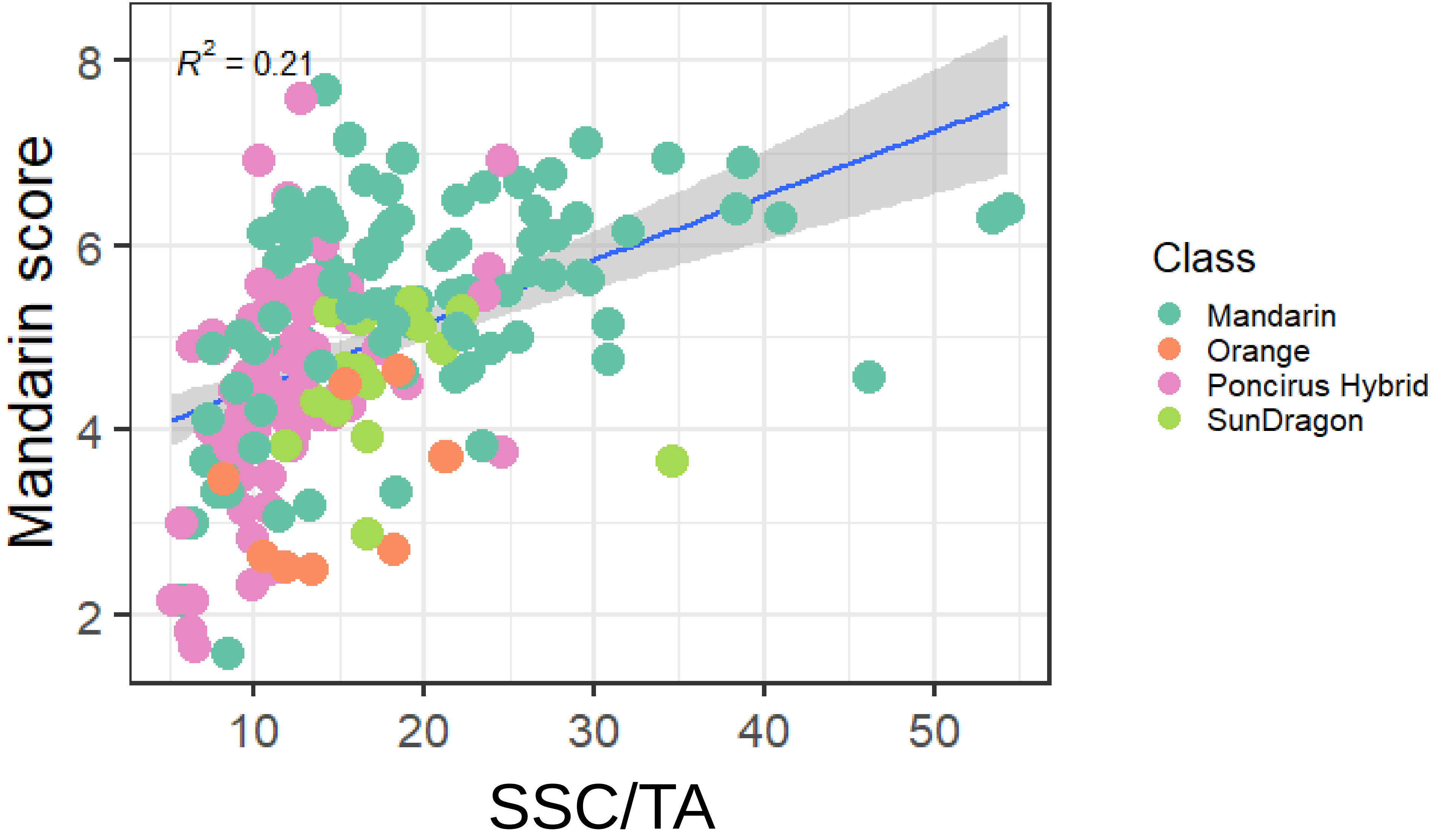


Figure S1. Mandarin flavor scores regressed against SSC/TA. Individual dots are colored according to their breeding class. The best fitted linear model is plotted with 0.95 confidence intervals. R squared value is annotated on the top left.

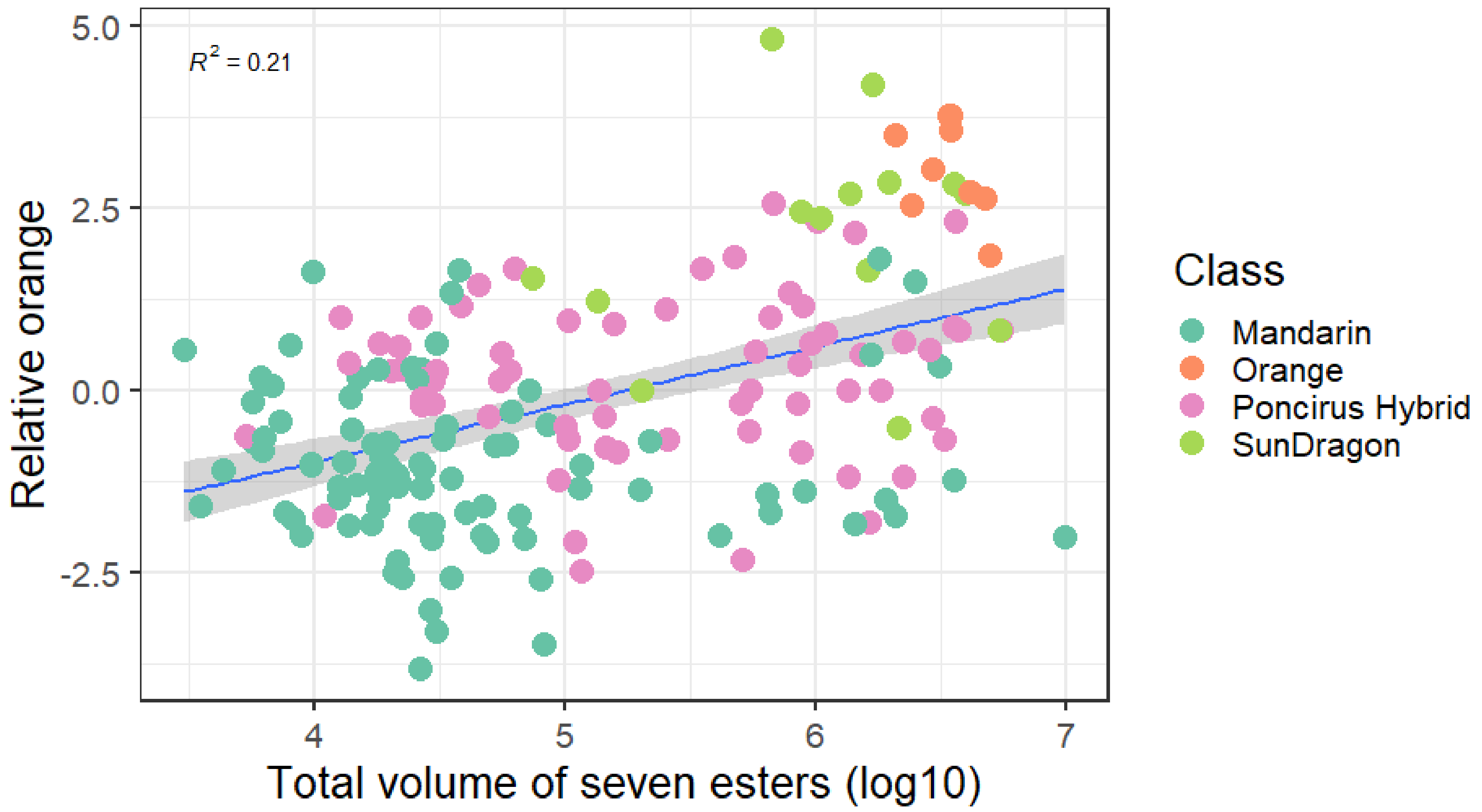


Figure S2. Relationship between relative orange flavor scores (orange flavor – mandarin flavor) and total volume of seven esters. Individual dots are colored according to their breeding class. The best fitted linear model is plotted with 0.95 confidence intervals. R squared value is annotated on the top left.

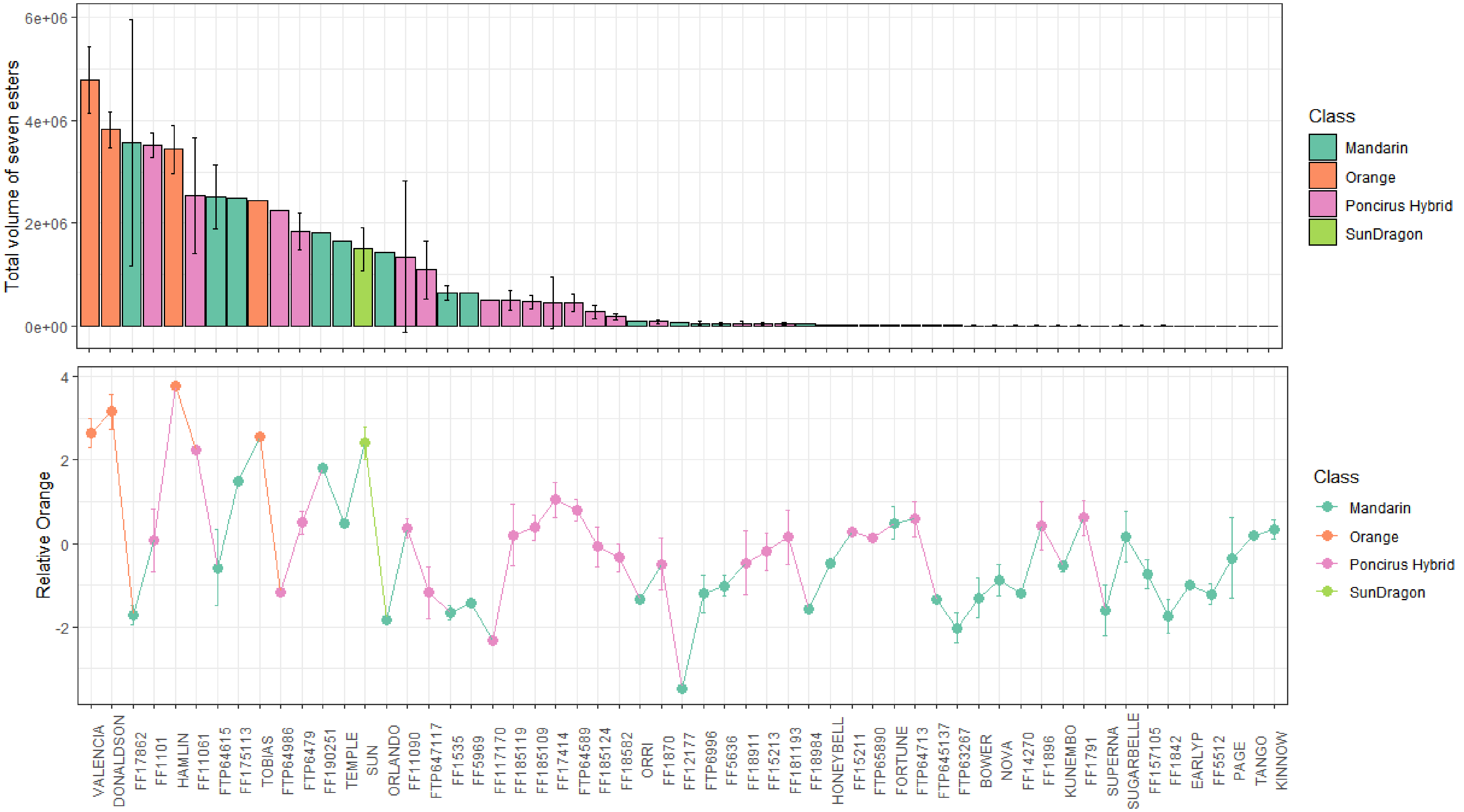


Figure S3. The total volume of seven esters (top) and relative orange score (bottom) are plotted for each genotype. Bars and dots are colored according to their breeding class. Error bars are calculated based on multiple harvests.

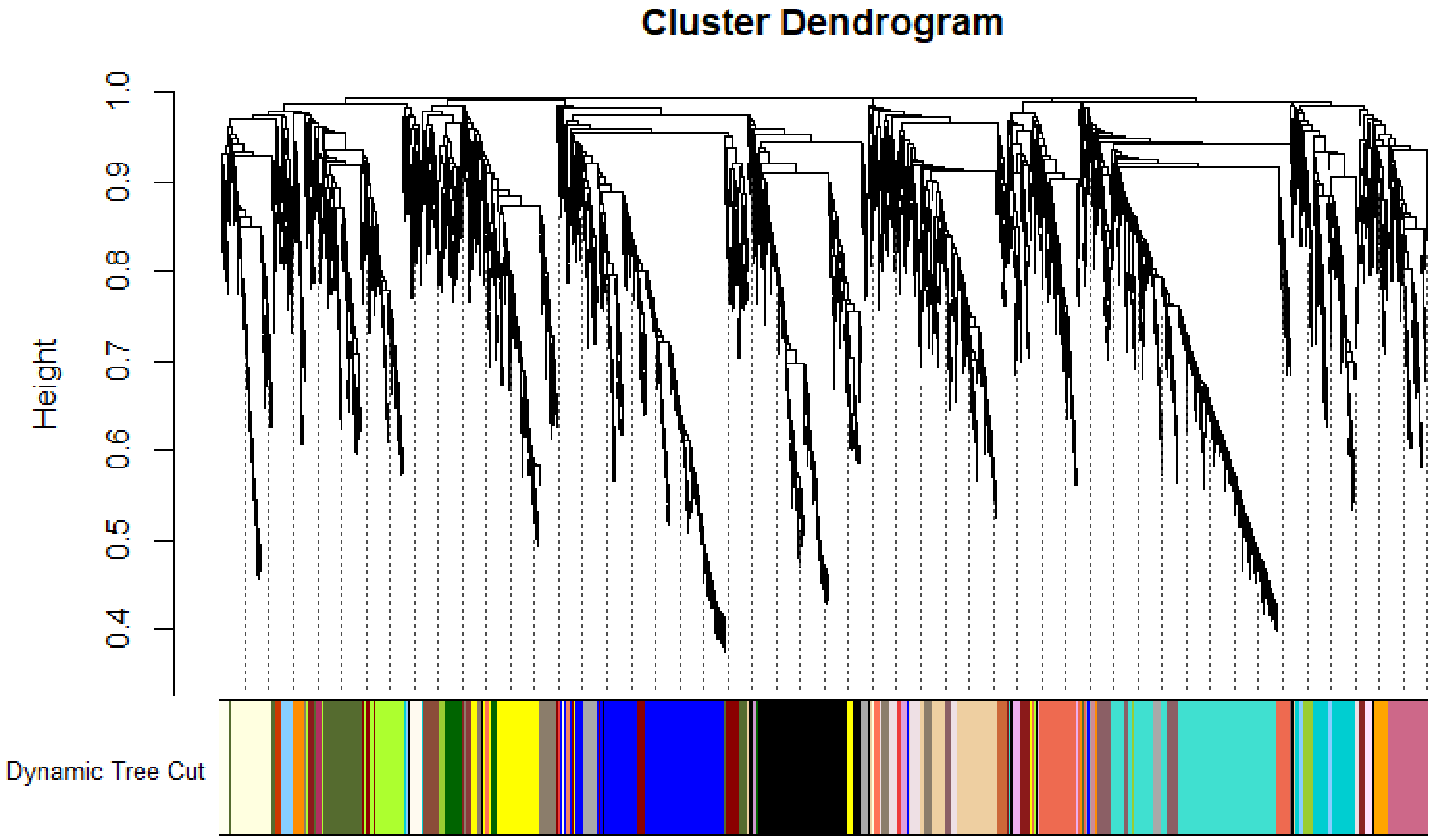


Figure S4. Co-expression modules assigned using weighted correlation network analysis (WGCNA). The top plot shows hierarchical clustering results. The bottom track shows the assigned module based on the dynamic tree cutting algorism.

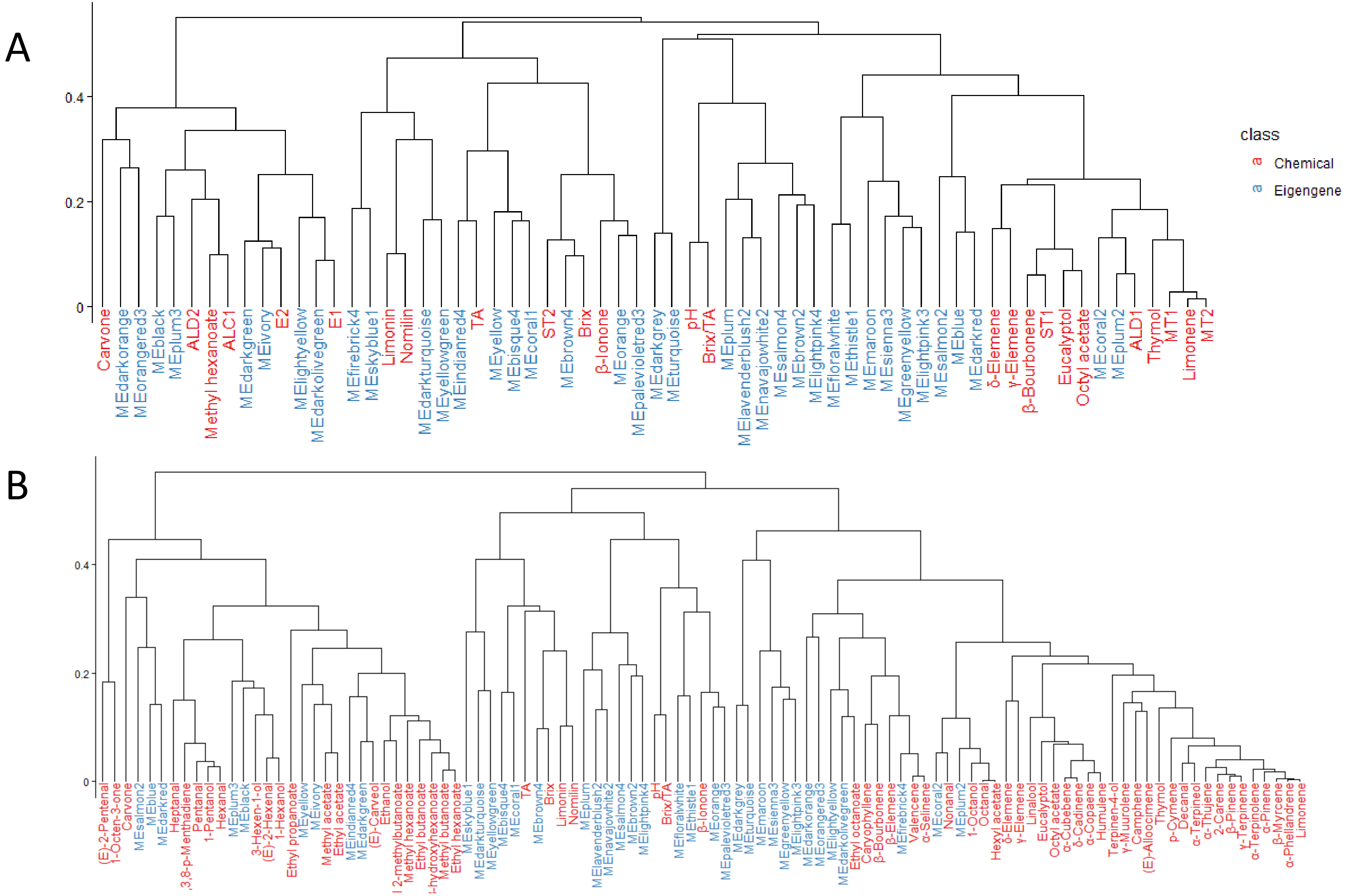


Figure S5. Dendrogram of correlations among eigengenes (the first eigenvector) of co-expression modules and the first eigenvectors of chemical clusters (A). Dendrogram of correlations among eigengenes of co-expression modules and individual compounds (B).

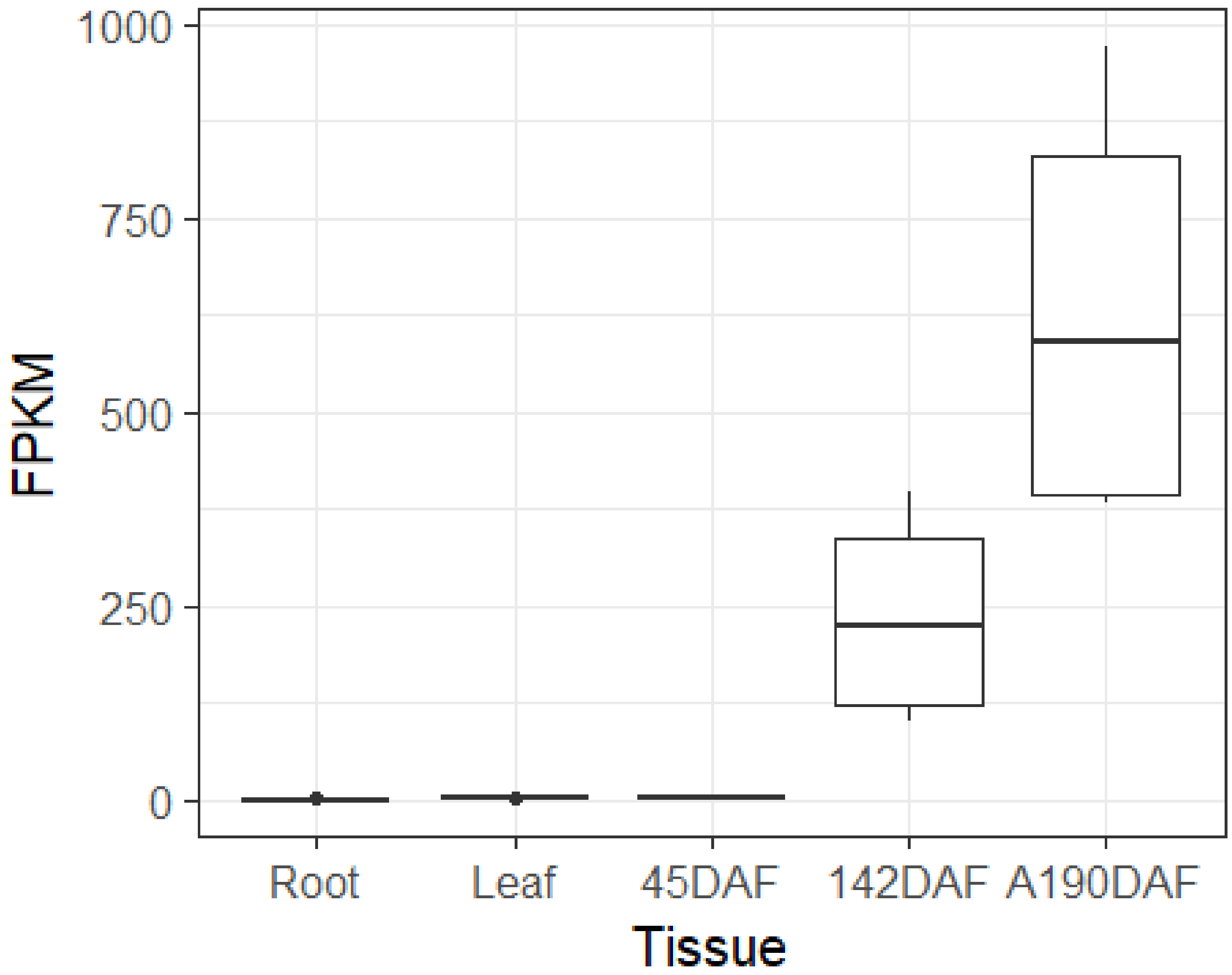


Figure S6. Expression of *CsAAT1* across five different tissues. 45DAF and 142DAF : fruits harvested 45 and 142 days after flowering, respectively. A190DAF: fruits harvested after 190 days after flowering. Raw data were downloaded from public RNAseq results deposited at CPBD (http://citrus.hzau.edu.cn/index.php). Four replicates were used for each tissue.

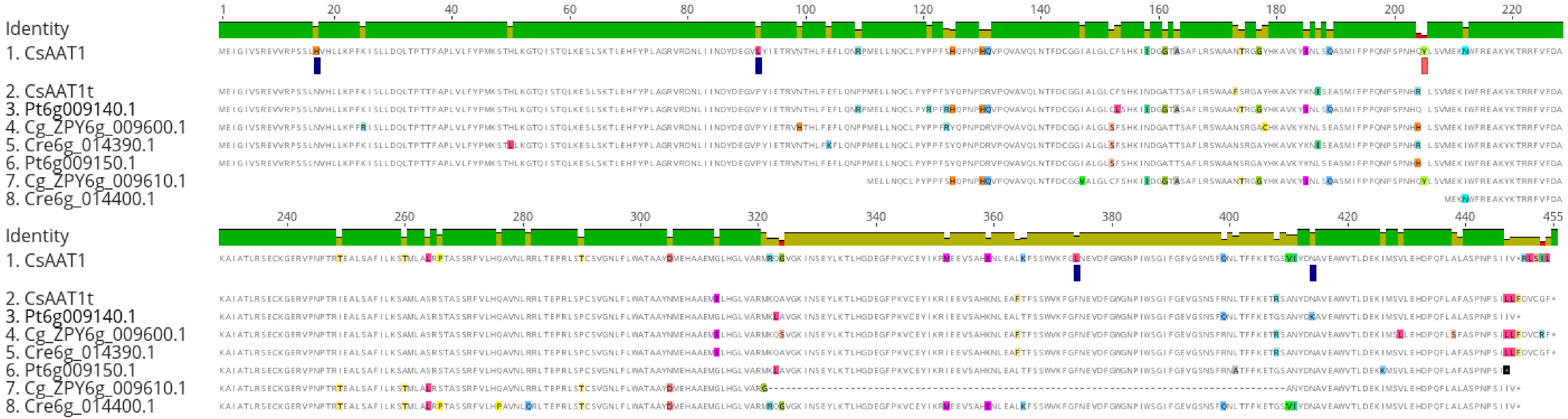


Figure S7. Protein alignment among homologs of *CsAAT1* in *Poncirus trifoliata* (Pt6g009140.1 & Pt6g009150.1 ), *Citrus reticulata* (Cre6g_014390.1 & Cre6g_014400.1) and *Citrus maxima* (Cg_ZPY6g_009600.1 & Cg_ZPY6g_009610.1) with *CsAAT1* and *CsAAT1t*.

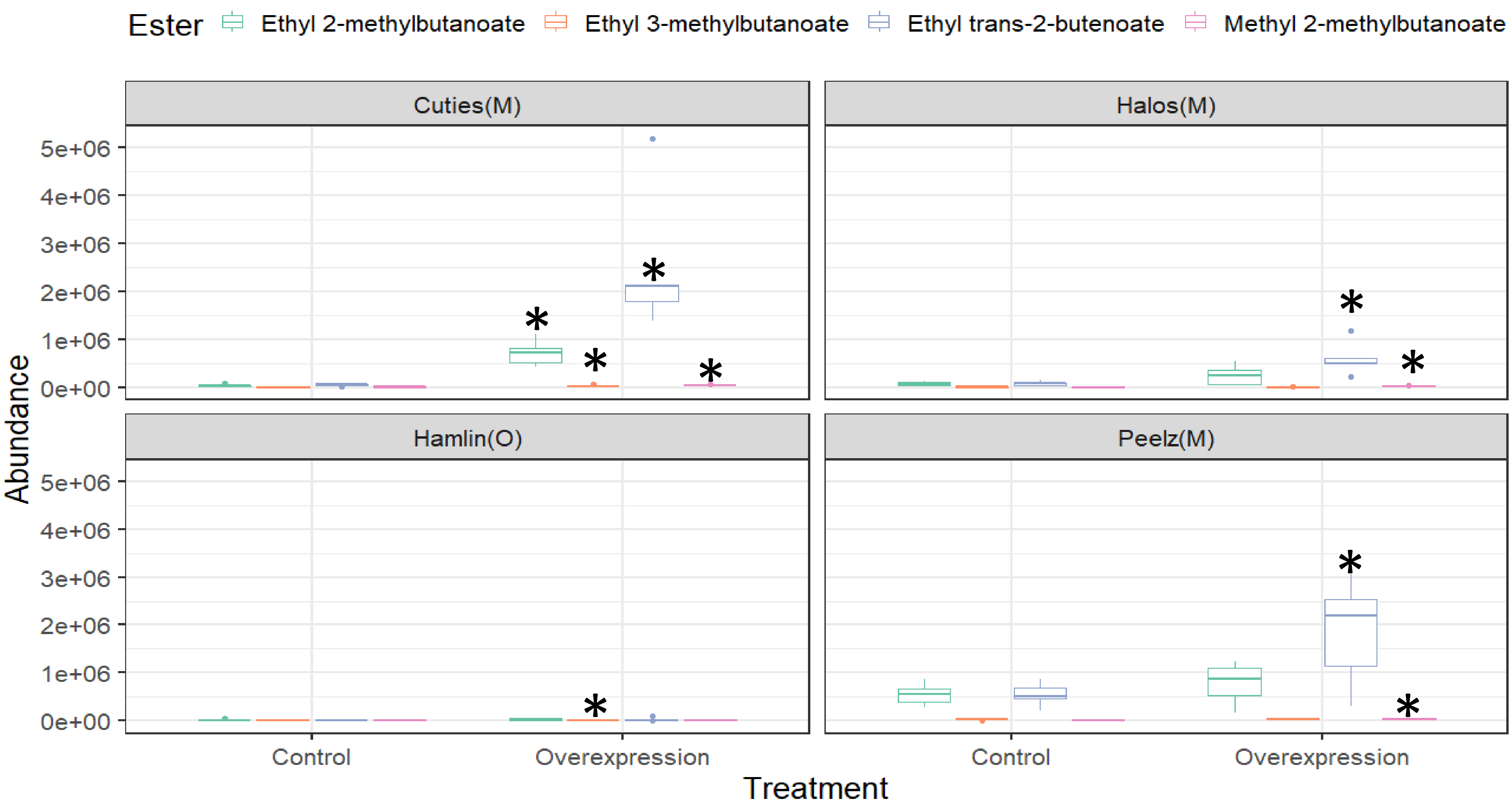


Figure S8. Comparisons of production for four branched-chain esters between control citrus fruits and fruits agroinfiltrated with *CsAAT1* overexpression construct. M indicates mandarin cultivar. O indicates orange cultivar. Asterisks indicate P-values < 0.05 based on Student t tests.

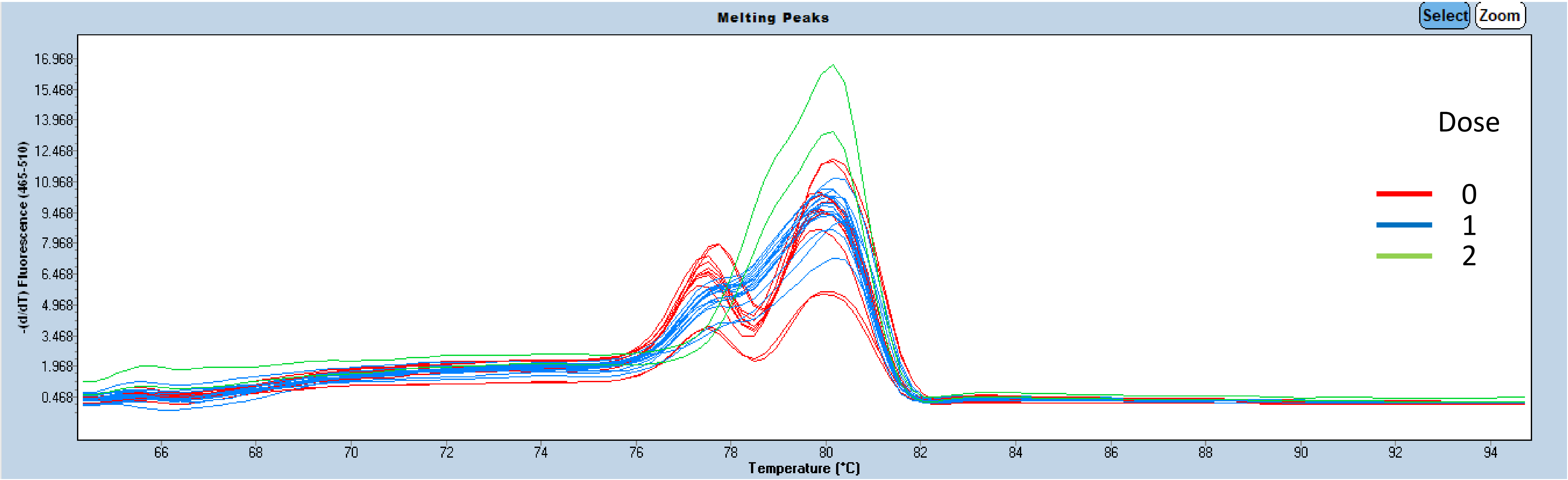


Figure S9. Melting curve patterns of samples with different dosages of orange allele of *CsAAT1.*
